# Beyond the Colony-Forming-Unit: Rapid Bacterial Evaluation in Osteomyelitis

**DOI:** 10.1101/2023.11.21.568051

**Authors:** Qi Sun, Kimberley Huynh, Dzenita Muratovic, Nicholas J. Gunn, Anja R. Zelmer, L. Bogdan Solomon, Gerald J. Atkins, Dongqing Yang

**Affiliations:** Centre for Orthopaedic & Trauma Research, Faculty of Health and Medical Sciences, University of Adelaide, Adelaide, SA, Australia; Department of Orthopaedics and Trauma, Royal Adelaide Hospital, Adelaide, SA, Australia

## Abstract

Examination of bacteria/host cell interactions is important for understanding the aetiology of many infectious diseases. The colony-forming-unit (CFU) has been the standard for quantifying bacterial burden for the past century, however, this suffers from low sensitivity and is dependent on bacterial culturability *in vitro*. Our data demonstrate the discrepancy between the CFU and bacterial genome copy number in an osteomyelitis-relevant co-culture system and we confirm diagnosis and quantify bacterial load in clinical bone specimens. This study provides an improved workflow for the quantification of bacterial burden in such cases.

---

Co-culturing mammalian host cells with bacteria *in vitro* is an important tool for examining the host-pathogen interaction in infectious disease modelling [1]. One of the essential readouts is the intracellular bacterial burden carried by the infected host cells. The most commonly used method for enumerating such bacterial load is counting the colony-forming-unit (CFU) number by agar plate culturing following the lysis of host cells and release of bacteria [2]. However, the reliability of such measurements is impacted by various factors, including, but not limited to, host cell/tissue type, bacterial strain, bacterial load, and culturing conditions for CFU enumeration [3, 4]. Osteomyelitis (OM), an infectious disease with pathogen-mediated infection in bone tissues, features a considerable proportion of infected but culture-negative cases, ranging between 20-40% [5-7]. This clear deficiency in CFU evaluation create difficulties for the accurate diagnosis and therefore treatment of OM.

*Staphylococci*, including coagulase-negative species, such as *S. epidermidis*, and *S. aureus*, comprise the major pathogens in adult OM [8]. In this study, we employed a previously established osteocyte-like cell model, differentiated SaOS2 (SaOS2-OY) and the single most common causative pathogen in OM, *S. aureus*, to simulate the *in vitro* infection of osteocytes, the most abundant cell type in bone tissue [9, 10]. Two previously characterised *S. aureus* strains, a high virulence (methicillin-resistant *S. aureus*; MRSA) strain, WCH-SK2 (SK2), shown previously to establish an intracellular infection in human osteocytes [11] and a low virulence (methicillin-sensitive *S. aureus*; MSSA) strain, WCH-SK3 (SK3) [12], were chosen for experimentation. In addition to CFU enumeration to quantify bacterial number, a polymerase chain reaction (PCR) approach measuring genomic DNA copies represented by a single copy gene within the *S. aureus* genome, was also performed in parallel. In the past, real time quantitative PCR (qPCR) assays have been widely used for the quantification of bacteria. However, the working principle of standard qPCR is dependent on the establishment of a standard curve using a known number of bacterial gene copies and the fitting of test samples into a linear regression to estimate the bacterial number for each assay [13, 14]. Here, we utilised digital droplet PCR (ddPCR) for this measurement. By its nature, the latter technique offers the advantage of counting the absolute target sequence copy number within the sample and removes the need for normalisation [15]. For all PCR-based assays, the quality of DNA containing the target sequence for amplification and quantification is critical for the sensitivity and reproducibility of readout. Therefore, methods for DNA preparation from both human and bacterial cells were examined. Here, we introduced the usage of DirectPCR™ Lysis Reagent (Direct buffer), a lysis buffer that maximises the release of genomic DNA from samples and is compatible with the downstream PCR analysis from raw cell culture lysate without any purification step. With the application of Direct buffer for DNA preparations in comparison to a commonly used DNA extraction spin column kit (DNA kit), the absolute bacterial genome copy number quantified by ddPCR was measured to be 5-fold higher in SaOS2-OY samples and 100-fold higher in samples from two strains of *S. aureus*, SK2 and SK3; better overall reproducibility among biological replicates was also achieved using Direct buffer over the standard approach (**Fig. 1A-C**). Serial 10-fold dilutions of bacterial log-phase suspension cultures were used to generate DNA lysates for testing the extent of DNA release. Plotting genome copy number against CFU from the same cultures showed near perfect linear regression relationships (**Fig. 1D-E**), confirming the complete and highly reproducible release of bacterial DNA. With this validating readout, we sought to determine if the elimination of purification steps by using the Direct buffer might minimise the sample loss associated with the utilisation of conventional column-based extraction methods [16]. Further, the release of genomic material by this reagent appeared to be complete and unbiased in our experiments, which led to higher consistency of inter- and intra-assays.

**Fig. 1:**
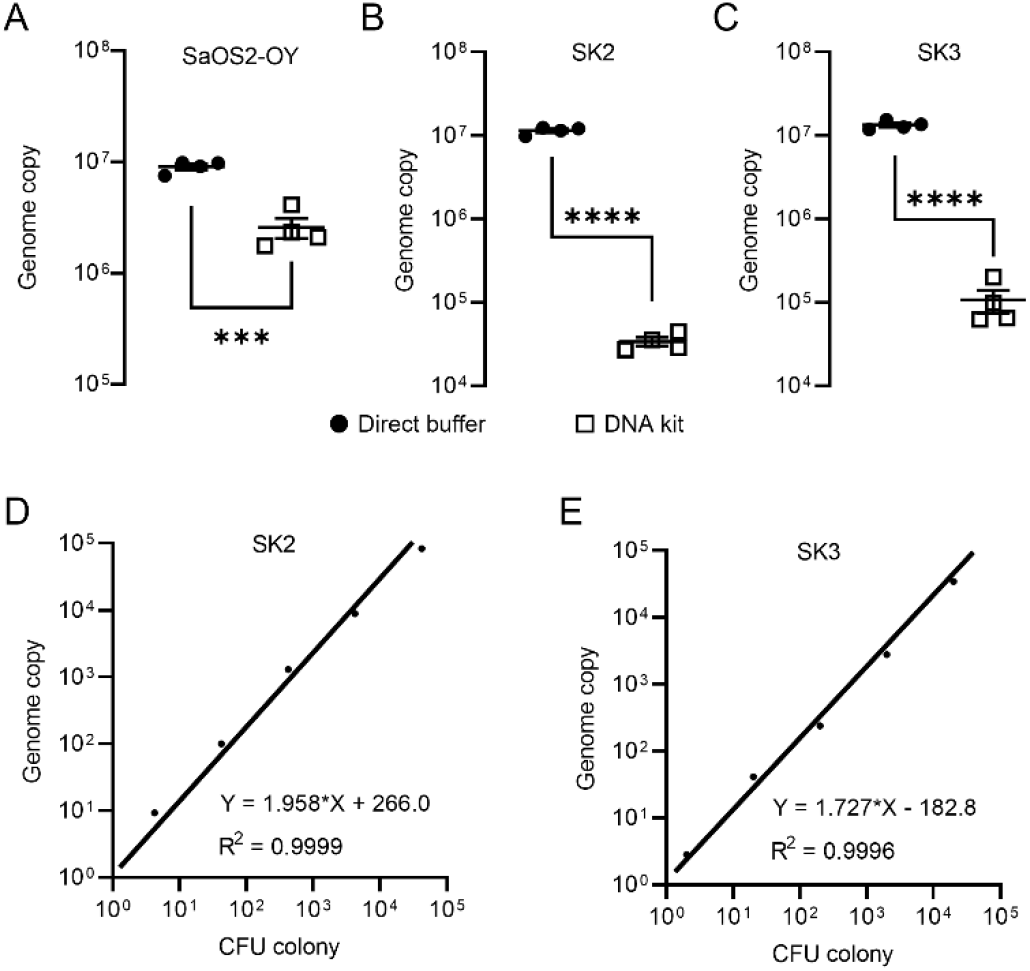
ddPCR genome counting of SaOS2-OY (A), SK2 (B) and SK3 (C), comparing the Direct buffer approach (•) and a standard DNA kit (□) (4 biological replicates with mean and standard errors were shown on graphs, ***p < 0.001 and ****p < 0.0001); demonstration of complete bacterial chromatin release from SK2 (D) and SK3 (E): correlation with CFU plating. Each data point represented the comparison of one CFU recovery and one total DNA measurement; three independent experiments were carried out for each strain and similar results were achieved; representative results of one experiment are presented.

Using the combined approach of direct lysis DNA preparation and ddPCR quantification, we then examined the *S. aureus* persistence in the intracellular environment in infected SaOS2-OY cells. Following infection, the SaOS2-OY and *S. aureus* co-cultures were lysed to break down host cell structures for the release of intracellular *S. aureus* for CFU development on agar plates. Consistent with previous findings, significant reductions ranging between 10^3^-fold and 10^6^-fold in cultured CFU on agar plates over the 5-day co-culture time period were observed in both SK2 and SK3 infected SaOS2-OY groups (**Fig. 2 A-F**). This trend was also found in low multiplicities of infection (MOI) (∼10^7^ total CFU/well, **Fig. 2 C & E**) and high MOI (∼10^8^ total CFU/well, **Fig. 2 D & F**) for initial bacteria-inoculated groups with either strain. However, when total DNA preparations from the whole culture lysates were taken for ddPCR quantification of bacterial genome copy number, differences in the dynamic changes of bacterial load were found. In the high virulence strain SK2-infected group, the genome copy numbers by 5 days post-infection were maintained at the levels of 10^7^ and 10^8^ cells/well for the low and high MOI groups, respectively (**Fig. 2 C & D**), similar to the original input levels.

**Fig. 2:**
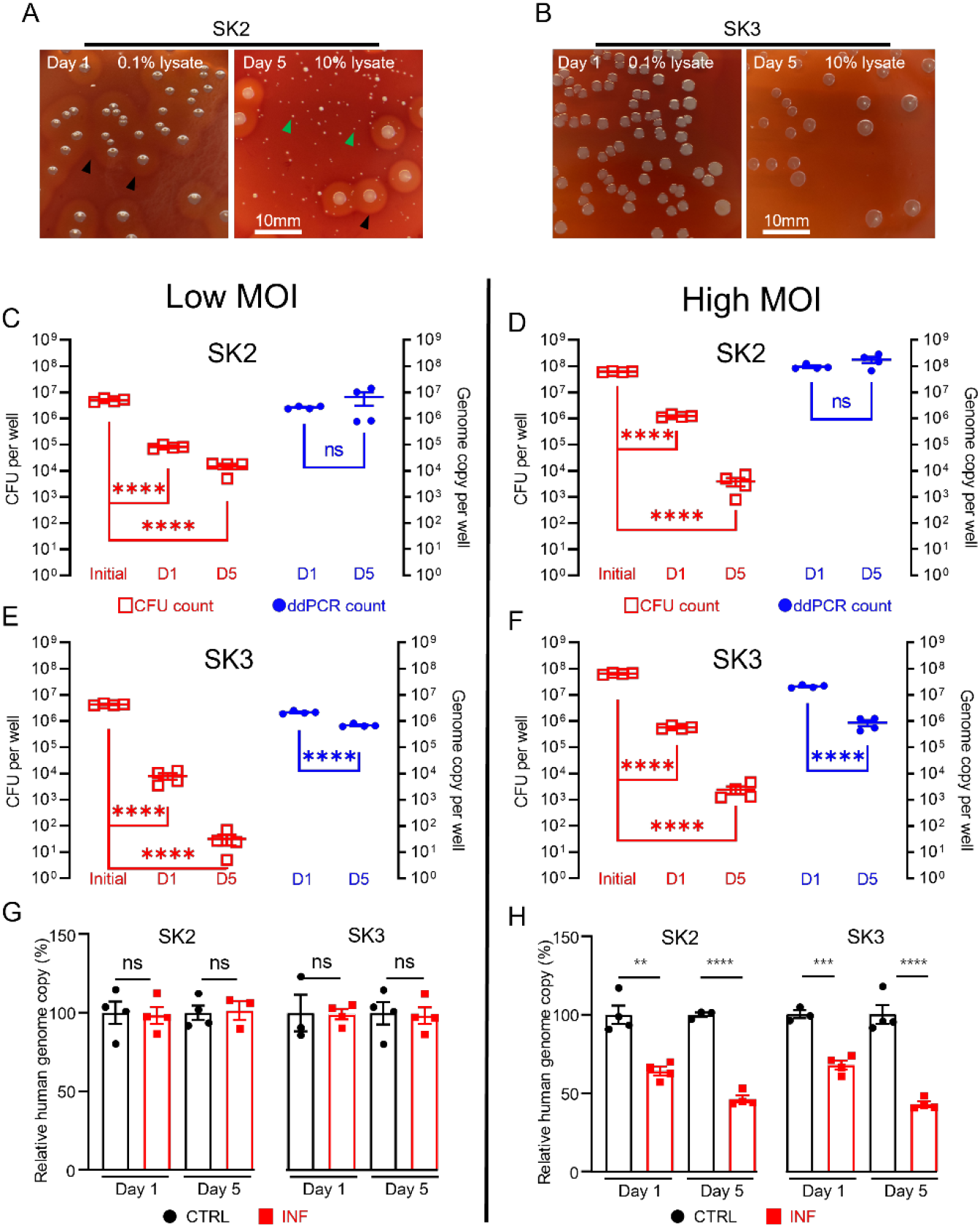
CFU recovery of SK2 (A) and SK3 (B) from host SaOS2-OY cells, with haemolysis reactions 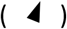 and numerous small colony variants 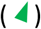 shown in SK2 group ; quantification of SK2 and SK3 using CFU count (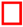, left Y) and ddPCR count (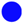, right Y) from low (C & E) and high (D & F) MOI groups; relative human genome copy (%) with CTRL (□) *VS* INF 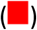 quantified by ddPCR from low (G) and high (H) MOI groups. (4 biological replicates with mean and standard errors are shown; ** p < 0.01, *** p < 0.001, **** p < 0.0001; ns = not significant)

However, the low virulence strain SK3-infected group demonstrated a decrease in genome copy counts. Interestingly, regardless of initial input amount, by 5 days post-infection, the remaining SK3 genome counts dropped to approximately 10^6^ cells/well for both groups (**Fig. 2 E & F**). The quantification of human genome copy number was also performed in all groups to estimate the remaining host cells in culture and this also served as an indicator of host cell viability. With the low MOI co-cultures, the human genome copy counts in the presence of either strain of *S. aureus* were not statistically different from uninfected controls, across 1- and 5-days post-infection. When the initial bacterial inoculum was increased 10-fold to 10^8^ cells per well, reductions in the human genome copies of ∼30% and 50% were measured in 1- and 5-day post-infection groups, respectively; such a trend was found to be similar in both SK2- and SK3-infected groups. Overall, our observations demonstrated profound differences in the readout of the presence of intracellular bacteria when using CFU and ddPCR quantification methods. Such inconsistency between these readouts in our model was as high as 10^6^-fold, with interaction of factors including initial bacterial load and strain variation. In contrast, comparison of the same quantification methods performed on bacteria suspension cultures yielded differences under 2-fold, indicated by the respective slope factor (less than 2) in the linear regression curves for both SK2 and SK3 strains (see **Fig. 1 D & E**). Together, our results indicated the dramatic change in bacterial culturability when comparing growth in ideal microbial suspension culture conditions and the growth limiting intracellular environment. Therefore, we propose that the interaction between the host cell response to infection and pathogen adaptation to the intracellular environment will compromise the reliability of the standard CFU counting method alone for the evaluation of bacterial persistence in a bacteria/host cell co-culture experiment. However, phenotypic variations were also observed in the recovered colonies from agar plating, with variations in haemolytic activity and small colony variant (SCV) formation, particularly in the SK2 group (**Fig. 2A**). SCV variants of *S. aureus* feature reduced metabolic activity and growth but are an indication of adaptation and the establishment of a chronic infection. This is consistent with the findings of reductions in CFU recovery but maintained levels of detectable genome copies, at least in the SK2-infected group.

To examine the applicability of this workflow to a clinical setting, we next extended the tests to clinical human bone specimens. Three clinically culture-negative periprosthetic joint infection (PJI) cases were chosen for analysis. All three cases were confirmed infected by clinical observation for typical PJI symptoms by clinicians, using the current gold-standard Musculoskeletal Infection Society (MSIS) criteria [17]; while each patient presented with a sinus tract communicating with their prosthesis, a major diagnostic criterion, neither the culturing of the synovial membrane biopsies nor synovial fluid around the prosthesis returned any positive cultures from a clinical laboratory. The bone tissue sections from the 3 patients were examined by Masson’s trichrome staining in comparison to tissue section from an osteoarthritis (OA) patient as non-infection control, to examine the bone matrix degradation, according to our previous study [18]. With this method, the red colour indicates intact bone matrix collagen and the blue colour corresponds to degraded collagen. Within all the histological observations, matrix degradation was found in sections of all three PJI patients (**Fig. 3 B-D**), whereas the staining of the control OA patient bone indicated intact collagen (**Fig. 3 A**), consistent with infection in the 3 PJI patient specimens [18]. In addition to the gold-standard diagnostic approach and histological staining, we performed CFU analysis using homogenised bone tissue, and negative culture results were also recorded. In this study, the samples containing total genomic DNA were prepared from histological sections in conjunction with the use of the DirectPCR™ Lysis reagent. The prepared DNA samples were put through a PCR amplification using primer set targeting a highly conserved bacterial genomic region, elongation factor Tu (*tuf*) [19], for the enrichment of bacterial signal within host/pathogen genomic mixture. The generated amplicons were then sequenced using an Oxford Nanopore Technology (ONT) Minion sequencer for the identification of bacterial species. All three patient bone DNA samples were confirmed to be bacterial genomic positive, with combinations identified of the coagulase-negative staphylococcal species *S. haemolyticus, S. hominis* and *S. epidermidis* (**Fig. 3 E-G**). The respective bacterial load for each of the samples was then quantified by ddPCR. Each bone sample examined contained between 2 × 10^4^ and 1 × 10^6^ bacterial genomic copies per million human genomic copies (**Fig. 3 E-G**). To confirm that the detected bacteria species were not introduced during the handling procedures, bone specimens from five primary total hip replacement (non-infected) patients were processed at the same time along with the PJI specimens; PCR reactions with negative *tuf* amplification for these bone samples indicated that operational contamination was unlikely (**Supplementary Fig. 1**).

**Fig. 3:**
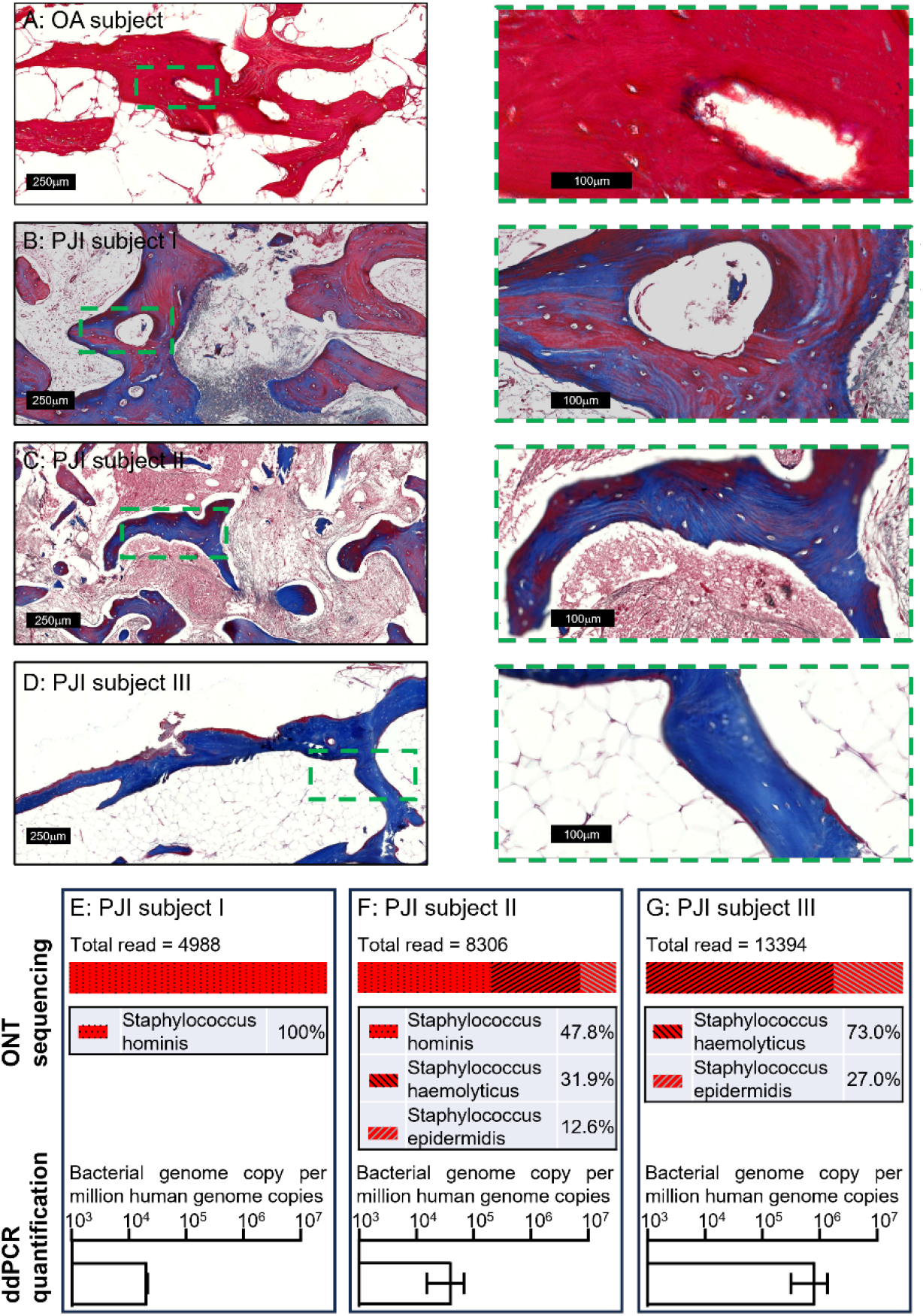
Masson’s trichrome staining on bone tissue sections of an osteoarthritis (OA) subject (A) and culture negative PJI subjects I-III (B-D). Pathogen profiling using total DNA of bone specimens from the above 3 PJI patients (E-G) by ONT sequencing for the readout of pathogen species and ddPCR to quantify bacterial load as a ratio of bacterial : human genomic copies (error bars shown are the combination of the standard error of the mean from two individual DNA preparations and the device generated error from each of the ddPCR runs, with over 15,000 droplets read per sample).

In this study, by the analysis of genomic material, we have addressed the ineffectiveness of the classical bacterial CFU development approach in determining and quantifying a confirmed infection by both an *in vitro* experimental model and by examining human clinical cases. With the introduction of a direct DNA release approach, less handling is required while delivering increased accuracy. The addition of ddPCR provided the advantage of achieving absolute quantification without the need for a PCR standard curve, which is both cost and time effective. For the purposes of unknown pathogen diagnosis in clinical cases, the exact bacterial species readout is required from sequencing the generated amplicons. Here, the utilisation of Oxford Nanopore technology dramatically lowered the equipment and skill demands for performing the analysis, since the MinION sequencer is a portable device and the sequencing results are analysed by the on-board software with little bioinformatics requirement. Noted here, the usual whole genome sequencing approach might not be suitable for this application, as in many of the clinical specimens, the amount of human genome can be orders of magnitude higher than the level of bacterial genomic material. Sequencing and analysing relatively rare bacterial DNA in this scenario is considerably more lengthy and costly. Therefore, a PCR approach was chosen to amplify the bacterial signal for detection. In this study, bacterial gene *tuf* was chosen as the target sequence to generate PCR amplicons instead of the commonly used *16S* rRNA gene sequence, because the prior one has been shown greater discriminatory power than *16S* [20]. Here, we took *tuf* sequences from *S. aureus* and the three coagulase-negative *staphylococcal* species identified in the clinical specimens for cross-comparisons (**Supplementary Fig. 2A**). Within the designated amplified sequence of only 271 bp, 11-22 (or 4.1%-8.1% differences) mismatches were identified between these four species (**Supplementary Fig. 2B**). For achieving such equivalent differences in *16S* sequences among the four strains, PCR amplicons with over 1000 bp in length are required (data not shown). Again, the strategy of analysing a short amplicon not only offers reduced reaction time, more importantly, it minimises the chance of generating non-specific products by shortening the elongation step. The listed primer set targeting the *tuf* gene was confirmed to cover most of the species in genus *Staphylococcus in silico*. In practice, it is challenging to find universal primers that cover the entire bacterial domain. Hence, for future real-world tests in clinical diagnosis, the strategy of using multiple primer sets to cover common PJI pathogens might need to be considered. Together, this workflow is potentially applicable as a point-of-care diagnostic method with the prospect of a rapid turnover: within hours comparing to the current procedure in the magnitude of days.

To conclude, for the purpose of quantifying the intracellular bacterial load in a host-pathogen co-culture setting, we have developed a workflow with the advantages of: a) rapid and labour-saving, with the elimination all the DNA extraction steps by virtue of using the direct lysis approach; b) minimisation of potential sample loss encountered with conventional DNA isolation methods, enabling better accuracy and reproducibility; c) absolute number quantification by ddPCR for achieving superior consistency of intra-assay experiments; d) sequencing analysis using the ONT MinION platform has the potential to become a point-of-care diagnostic approach when performing unknown pathogen identification in a clinical setting. Nevertheless, we suggest that the CFU plating method should not be omitted as it allows the evaluation of bacterial phenotypic adaptation in such experimentation.

## Materials and methods

### SaOS2 cell culture and differentiation

The human osteosarcoma cell line SaOS2 differentiated to an osteocyte-like stage was employed for performing the bacteria/host co-culture experiments, as previously described [10]. Cells were seeded at a density of 2 × 10^4^ cells per well in 48-well plates and cultured at 37°C/5% CO_2_, in growth media consisting of αMEM (Thermo-Fisher, Victoria, Australia) supplemented with 10% v/v foetal calf serum (FCS) (Thermo-Fisher), 2 mM L-Glutamine (Thermo-Fisher), 1 U/ml penicillin/streptomycin (Thermo-Fisher). At 90% confluence, SaOS2 cultures were switched to osteogenic differentiation medium comprising aMEM supplemented with 5% v/v FCS, 100 mM ascorbate 2-phosphate (Sigma-Aldrich, St Louis, USA), 1.8 mM potassium di-hydrogen phosphate (Sigma-Aldrich), 1 mM HEPES (Thermo-Fisher), 2 mM L-Glutamine and 1 U/ml penicillin/streptomycin. Cells were then maintained under differentiation conditions for 28 days to achieve osteocyte-like phenotype (SaOS2-OY) [9].

### *S. aureus* preparation

Two pre-characterised *S. aureus* strains, an MRSA strain, WCH-SK2 (SK2) [12] and an MSSA strain, WCH-SK3 (SK3), were grown in Terrific Broth (Thermo-Fisher) on a 37°C /200 rpm rocking platform to achieve log-phase suspension cultures individually. Bacteria were pelleted by 3000 g for 5 min centrifugation and then resuspended in sterile phosphate-buffered saline (PBS) to estimate cell number by the optical density (OD) at 600 nm light absorption. The two bacteria suspensions were then taken to undergo 10-fold serial dilutions and the diluents were individually plated on blood agar plates containing 10% v/v defibrinated sheep blood (Thermo-Fisher) to determine the colony-forming-unit (CFU) number from original bacterial suspension preparations. The calculated CFU numbers from agar plates were compared to the ddPCR quantified genome copy numbers.

### Host cell infection by co-culturing of SaOS2-OY and *S. aureus*

For host cell infection, bacterial suspension cultures were resuspended in sterile PBS, as described above, to achieve the density for low and high multiplicities of infection (MOI). The differentiation media for SaOS2-OY was removed and the cells washed twice with PBS. The resulting bacterial inoculums were added to SaOS2-OY cells to allow invasion for 1 hour at 37°C/5% CO_2_. Post-infection, cultures were washed twice with PBS and incubated at 37°C/5% CO_2_with 20 µg/ml lysostaphin (Sigma-Aldrich) in antibiotic-free media for 24 h to eliminate extracellular bacteria before replenishment with fresh differentiation medium. Supernatants were streaked on agar plates to verify the absence of extracellular bacteria at 24 h and 120 h post-infection.

### Measurements of intracellular bacterial number by CFU counting

The intracellular bacteria numbers were determined at 24 h and 120 h post-infection time points by spreading serial dilutions of cell lysates on blood agar plates and culturing at 37°C /5% CO_2_for 48 hours.

### DNA extraction from host cells and bacteria

For genomic quantification, total DNA was isolated using either DNeasy Blood & Tissue Kits (Qiagen Inc., VIC, Australia), or DirectPCR™ Lysis Reagent (Direct buffer) (Viagen Biotech Inc., CA, USA), as per the manufacturers’ instructions. For the new approach of using the Direct buffer, 500 ml buffer together with 200 μg/ml proteinase K (Thermo-Fisher) was used for the lysis of one SaOS2-OY sample in one well from a 48-well tissue culture plate or one pelleted bacterial sample prepared from serial dilutions of a suspension culture. Individual cell lysates were transferred to 1.5 ml reaction tubes, digested at 55°C for 35 min and then heat-inactivated at 85°C for 15 min to terminate enzymic digestion. The processed lysates were then ready for PCR analyses.

### Digital droplet polymerase chain reaction (ddPCR)

The ddPCR assay was performed using QX200 Digital Droplet PCR System (Bio-Rad Laboratories, USA), as per the manufacturer’s instructions. Primer sets targeting a human genome-specific sequence within the single-copy type X collagen (*COL10A1*) gene and a *S. aureus* genome-specific sequence within the sigma factor B (*sigB*) gene, were used. The sequences of primer sets were:

*COL10A1*: forward 5’-ccaccaggtcaagcagtcat-3’, reverse 5’-gttggcactaacaagaggggt-3’

*sigB*: forward 5’-ggggcaacaagatgaccatt-3’, reverse 5’-tgccgttctctgaagtcgtg-3’

### Analysis of human bone tissues

All human studies received institutional research ethics approval (Royal Adelaide Hospital Human Research Ethics Committee Approval No. 14466). Bone biopsies were collected from patients undergoing either primary total hip replacement or revision total hip replacement surgery for PJI, with informed, written patient consent. Each bone specimen was separated into two parts. One of these was homogenised in a bead-beating tissue homogeniser (Bead Ruptor Elite, Omni International, Kennesaw, GA, USA), with 2 cycles of beating at 3 m/s for 30 s to disassociate the bone structure for the release of potentially viable bacteria. Bone homogenates were then sent to a clinical pathology laboratory (SA Pathology, Adelaide, Australia) for CFU analysis. Clinical soft tissue and fluid samples were sent to SA Pathology directly from surgery for pathogen analysis. The second piece of bone biopsy was processed and de-mineralised using OSTEOSOFT® (Sigma-Aldrich) solution for standard paraffin embedding procedure. Post-embedding, the bone tissues were sectioned under DNase/RNase-free conditions. One 5μm section from each bone specimen was used for Masson’s Trichrome staining, as described in our previous study [18], and six sections were pooled together in DirectPCR™ Lysis Reagent for the isolation of total genomic DNA, as described above. Prepared DNA samples were then subjected to 1) conventional PCR amplification for an ONT sequencing readout (MinION sequencer, Oxford Nanopore Technology, Oxford, UK), according to the manufacturer’s instructions; 2) ddPCR for the absolute quantification for both human and bacterial genome copy number. The primer sets targeting the human *COL10A1* genomic sequence, as listed above, and bacterial *tuf* sequence primers (forward 5’-ttctcaatcactggtcgtgg-3’, reverse 5’-ggagtatgacgtccaccttc-3’) were used for measuring human and bacterial genome copies, respectively. The output sequencing data were analysed by Epi2ME software within the WIMP workflow (Oxford Nanopore Technologies) for bacterial taxonomical analysis.

### Statistics

The results were graphed as means ± standard errors of the means (SEMs). The significance between two treatment groups was evaluated using two-tailed T-test using GraphPad Prism 10.2 (GraphPad Software, MA, USA), where applicable.

## Acknowledgement

This work was supported by a National Health and Medical Research Council of Australia (NHMRC) Ideas Grant scheme (ID 2011042) awarded to G.J.A. and D.Y. and an Australian Orthopaedic Association Research Grant awarded to G.J.A. and L.B.S.. Oxford Nanopore sequencing was supported by a University of Adelaide Faculty of Health and Medical Sciences infrastructure grant with technical assistance provided by Ms Thessa Kroes and Dr. Mark Corbett. Q.S. was supported by a University of Adelaide Faculty of Health and Medical Sciences Postgraduate Research Scholarship.

## Author contributions

D.Y., G.J.A. and L.B.S. conceived the study. D.Y. adapted key protocols and developed the workflow; Q.S. performed most of the experimental work; K.H. contributed to molecular analysis and D.M. contributed to histological analysis; N.J.G and A.R.Z validated the methods developed in this work; Q.S. and D.Y. drafted the manuscript; G.J.A and D.Y provided scientific insight and supervised the study; L.B.S. consented patients and performed biopsy retrieval; all authors contributed to manuscript preparation and data presentation and agreed on the submitted version of the manuscript.

---

**Supplementary figure 1:**
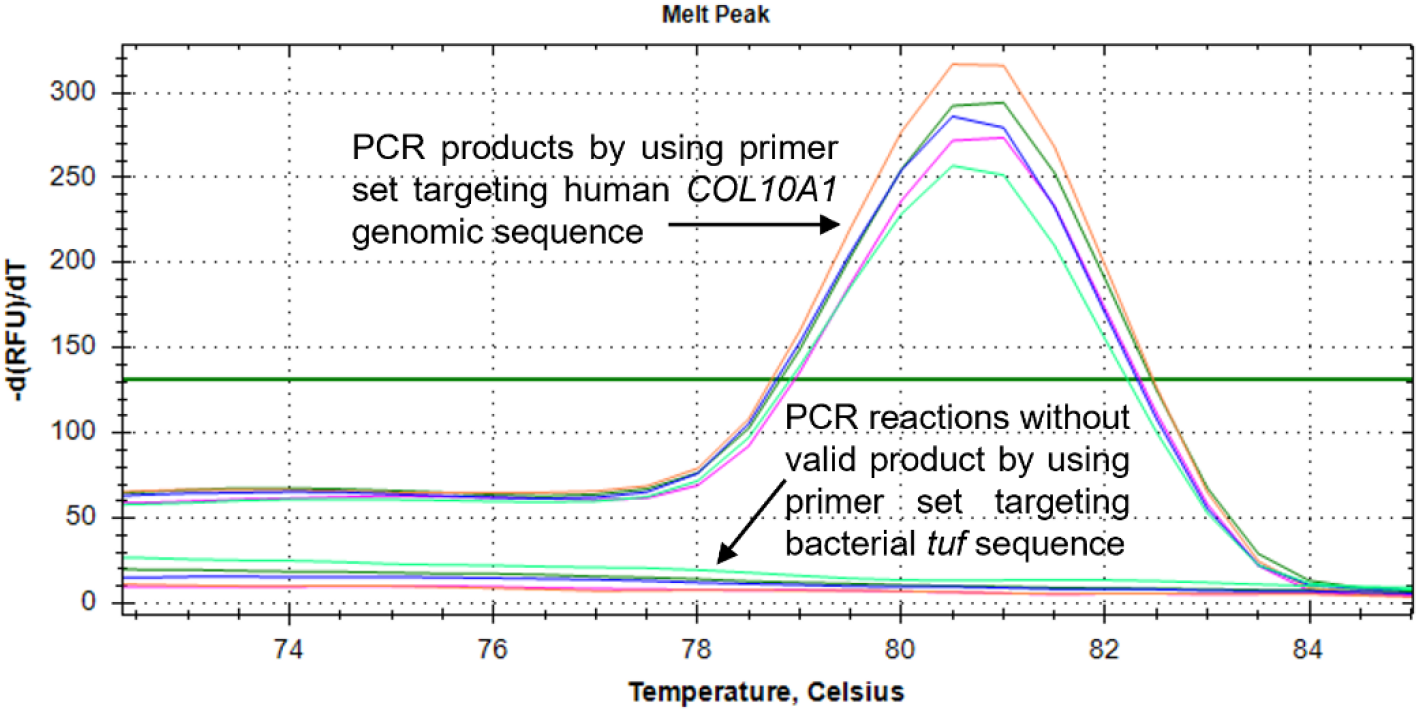
PCR analysis of bone samples from primary total hip replacement cases. DNA isolated from 5 patient bone samples were analysed by PCR for the presence of human *COL10A1* and bacterial *tuf*. The negative presence of bacterial *tuf* PCR product was confirmed by melt analysis post-quantitative PCR reactions using the DNA samples of five primary total hip replacement (non-infected) patients (coded with five different colours).

**Supplementary figure 2:**
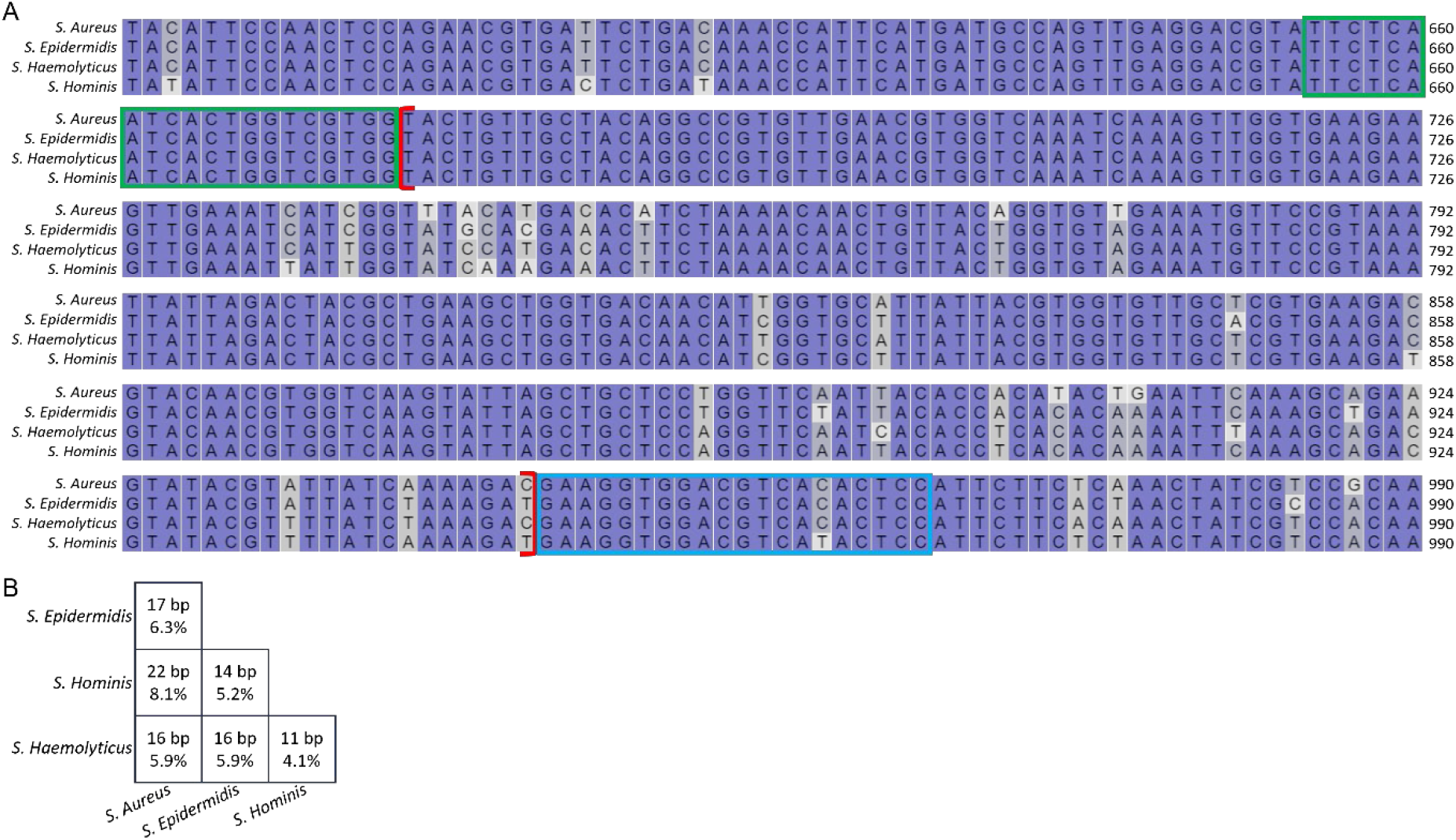
Multi-sequence alignment among *S. aureus, S. epidermidis, S. haemolyticus* and *S. hominis*. The green and blue boxes indicate the forward and reverse primer binding areas, respectively; and the region between the red brackets was the assaying sequence for analysis (A). A summary of the number of mismatched base pairs and % differences within the assayed regions, among the above four species (B).

## References

1. Hofstee, M.I., et al., Current Concepts of Osteomyelitis: From Pathologic Mechanisms to Advanced Research Methods. Am J Pathol, 2020. 190(6): p. 1151–1163.

2. Staley, J.T. and A. Konopka, Measurement of in situ activities of nonphotosynthetic microorganisms in aquatic and terrestrial habitats. Annu Rev Microbiol, 1985. 39: p. 321–46.

3. Chang, C.W., et al., Factors affecting microbiological colony count accuracy for bioaerosol sampling and analysis. Am Ind Hyg Assoc J, 1995. 56(10): p. 979–86.

4. Davis, K.E., S.J. Joseph, and P.H. Janssen, Effects of growth medium, inoculum size, and incubation time on culturability and isolation of soil bacteria. Appl Environ Microbiol, 2005. 71(2): p. 826–34.

5. Bejon, P., et al., Two-stage revision for prosthetic joint infection: predictors of outcome and the role of reimplantation microbiology. J Antimicrob Chemother, 2010. 65(3): p. 569–75.

6. Yoon, H.K., et al., A Review of the Literature on Culture-Negative Periprosthetic Joint Infection: Epidemiology, Diagnosis and Treatment. Knee Surg Relat Res, 2017. 29(3): p. 155–164.

7. Kim, Y.H., et al., Comparison of infection control rates and clinical outcomes in culture-positive and culture-negative infected total-knee arthroplasty. J Orthop, 2015. 12(Suppl 1): p. S37–43.

8. Kavanagh, N., et al., Staphylococcal Osteomyelitis: Disease Progression, Treatment Challenges, and Future Directions. Clin Microbiol Rev, 2018. 31(2).

9. Prideaux, M., et al., SaOS2 Osteosarcoma cells as an in vitro model for studying the transition of human osteoblasts to osteocytes. Calcif Tissue Int, 2014. 95(2): p. 183–93.

10. Gunn, N.J., et al., A Human Osteocyte Cell Line Model for Studying Staphylococcus aureus Persistence in Osteomyelitis. Front Cell Infect Microbiol, 2021. 11: p. 781022.

11. Yang, D., et al., Novel Insights into Staphylococcus aureus Deep Bone Infections: the Involvement of Osteocytes. mBio, 2018. 9(2).

12. Bui, L.M.G. and S.P. Kidd, A full genomic characterization of the development of a stable Small Colony Variant cell-type by a clinical Staphylococcus aureus strain. Infect Genet Evol, 2015. 36: p. 345–355.

13. Kralik, P. and M. Ricchi, A Basic Guide to Real Time PCR in Microbial Diagnostics: Definitions, Parameters, and Everything. Front Microbiol, 2017. 8: p. 108.

14. Smith, C.J. and A.M. Osborn, Advantages and limitations of quantitative PCR (Q-PCR)-based approaches in microbial ecology. FEMS Microbiol Ecol, 2009. 67(1): p. 6–20.

15. Hindson, C.M., et al., Absolute quantification by droplet digital PCR versus analog real-time PCR. Nat Methods, 2013. 10(10): p. 1003–5.

16. Schmitz, T.C., et al., Solid-phase silica-based extraction leads to underestimation of residual DNA in decellularized tissues. Xenotransplantation, 2021. 28(1): p. e12643.

17. Parvizi, J., et al., The 2018 Definition of Periprosthetic Hip and Knee Infection: An Evidence-Based and Validated Criteria. J Arthroplasty, 2018. 33(5): p. 1309–1314.e2.

18. Ormsby, R.T., et al., Evidence for osteocyte-mediated bone-matrix degradation associated with periprosthetic joint infection (PJI). Eur Cell Mater, 2021. 42: p. 264–280.

19. Li, X., et al., Use of tuf as a target for sequence-based identification of Gram-positive cocci of the genus Enterococcus, Streptococcus, coagulase-negative Staphylococcus, and Lactococcus. Ann Clin Microbiol Antimicrob, 2012. 11: p. 31.

20. Hwang, S.M., et al., Tuf gene sequence analysis has greater discriminatory power than 16S rRNA sequence analysis in identification of clinical isolates of coagulase-negative staphylococci. J Clin Microbiol, 2011. 49(12): p. 4142–9.

